# 4DPhenoMVS: A Low-Cost 3D Tomato Phenotyping Pipeline Using a 3D Reconstruction Point Cloud Based on Multiview Images

**DOI:** 10.1101/2021.11.09.467984

**Authors:** Ruifang Zhai, Yinghua Wang, Songtao Hu, Wanneng Yang

**Affiliations:** National Key Laboratory of Crop Genetic Improvement, National Center of Plant Gene Research (Wuhan), College of Informatics, Huazhong Agricultural University, Wuhan 430070, PR China

**Keywords:** dynamic 3D phenotyping, 3D reconstruction point cloud, structure from motion, growth analysis, whole growth stages, tomato

## Abstract

Manual phenotyping of tomato plants is time consuming and labor intensive. Due to the lack of low-cost and open-access 3D phenotyping tools, the dynamic 3D growth of tomato plants during all growth stages has not been fully explored. In this study, based on the 3D structural data points generated by employing structures from motion algorithms on multiple-view images, we proposed a dynamic 3D phenotyping pipeline, 4DPhenoMVS, to calculate and analyze 14 phenotypic traits of tomato plants covering the whole life cycle. The results showed that the R^2^ values between the phenotypic traits and the manual measurements stem length, plant height, and internode length were more than 0.8. In addition, to investigate the environmental influence on tomato plant growth and yield in the greenhouse, eight tomato plants were chosen and phenotyped during 7 growth stages according to different light intensities, temperatures, and humidities. The results showed that stronger light intensity and moderate temperature and humidity contribute to a higher growth rate and higher yield. In conclusion, we developed a low-cost and open-access 3D phenotyping pipeline for tomato plants, which will benefit tomato breeding, cultivation research, and functional genomics in the future.

**Highlights:** Based on the 3D structural data points generated by employing structures from motion algorithms on multiple-view images, we developed a low-cost and open-access 3D phenotyping tool for tomato plants during all growth stages.

## Introduction

The tomato, as the most popular vegetable crop, is widely cultivated worldwide under outdoor and indoor conditions due to its high nutrition and health benefits (Damtew 2017). It is essential to estimate the phenotypic traits of tomato plants and explore phenotypic trait variation during different growth stages. Although great advances have been made in tomato breeding, more efforts should be made to characterize objective, reliable and informative measurements of phenotypic traits to push breeding further (Daniel *et al.*, 2016). Thus, sustainable breakthrough in tomato phenotyping is urgently needed.

In the past decade, various phenotyping technologies have drawn much attention in the agricultural field because of the rapid development of new sensors and correspondingly high automation technology in the urgent need of non-destructivity (Jin *et al.*, 2020). Two-dimensional imaging technologies have been applied in structural trait estimation, growth analysis, and yield estimation at different time points (An *et al.*, 2017; Mu *et al.*, 2020; Yamamoto *et al.*, 2016; Zhang *et al.*, 2018). Although many image processing and analysis algorithms have been developed on various crops, several defects cannot be avoided, such as the ambiguity of plant size caused by camera viewpoints and camera-object distance, the lack of 3D information regarding plant volume, and self-occlusion problems caused by the complex structure of plants (Zhang *et al.*, 2018).

Thus, 3D phenotypic techniques have gradually become a powerful tool for obtaining phenotypic traits due to their noninvasive and noncontact properties and in particular their advantages in obtaining 3D geometric structural information compared to conventional 2D technologies (Zhang *et al.*, 2018). The superiority of using 3D information in the calculation of phenotypic traits, such as plant volume, plant height, and leaf length, has been demonstrated (Rose *et al.*, 2015). As 3D point clouds are becoming a standard data type for digitizing plant architecture in the laboratory or in the field, generating 3D points of crop plants is the essential challenge encountered in 3D phenotyping that needs to be addressed first. The most commonly used technologies for generating 3D points can be divided into two categories, namely, passive and active solutions (Wang *et al.*, 2018). For the last decade, 3D point clouds of plants have been able to be created by active sensors, such as in laser scanning. Laser scanning often outputs a complete and accurate 3D point cloud of one plant or block area of plants (Jin *et al.*, 2019); however, the high cost of laser scanners is the main bottleneck that limits their wide application. Compared to laser scanning technologies, passive reconstruction methods, usually camera-based, are still the most competitive and widely applicable technologies for reconstruction solutions because only an easily accessible and affordable camera is needed.

Generating the 3D structure of objects from a series of 2-D images is the first task that needs to be addressed before extracting 3D phenotypic traits. Multiview stereo (MVS) technology is the mainstream algorithm. Compared to binocular stereo vision, multiview stereo technology is more applicable because a complete point cloud of an object can be generated (Guo and Xu, 2017). Data collection can be completed in two scenarios: a fixed-camera scenario, in which one or multiple cameras are fixed in the scene, and a moving-camera scenario, in which a camera moves around the object in the scene. In the fixed-camera scenario, the camera remains in one location, and auxiliary equipment, such as a turn stand, is needed to take multiple images of one object from different viewpoints. Alternatively, multiple cameras can be equipped at different viewpoints with respect to the object in order to create a complete representation of the object. The earlier reconstruction process for plants usually used this strategy. Calibration is the most important prerequisite work, followed by different reconstruction strategies, mainly carving-based methods (Artzet *et al.*, 2019; Chen Yu, 2011; Das Choudhury *et al.*, 2020; Gibbs *et al.*, 2018; Golbach *et al.*, 2015; Roussel *et al.*, 2016; Ward *et al.*, 2014; Ying *et al.*, 2011) and correspondence-based methods (An *et al.*, 2017; Guo and Xu, 2017; Liu *et al.*, 2017; Lu *et al.*, 2020; Pound *et al.*, 2014; Wang *et al.*, 2018; Wu *et al.*, 2020; Yang and Han, 2020; Zhu *et al.*, 2020).

In the moving-camera scenario, a camera is moved around an object to capture multiple images from different viewpoints. In this case, structure from motion (SfM) and MVS are combined to generate a dense point cloud for objects, such as Arabidopsis (Liu *et al.*, 2017), tomato plants (Guo and Xu, 2017), Scindapsus and Pachira macrocarpa (Guo and Xu, 2017), maize plants (Wang *et al.*, 2018; Wu *et al.*, 2020), sweet potato plants (Zhang *et al.*, 2018), and even plant roots (Masuda, 2019).

Phenotyping methods of tomatoes have also been studied in several papers. In the conventional 2D imaging scenario, image analysis was mainly conducted on fruits and seeds (Daniel *et al.*, 2016). In addition, unmanned aerial vehicle (UAV)-based imagery and random forests were used for biomass and yield prediction at the field scale (Johansen *et al.*, 2020). Regarding the 3D reconstruction of tomato plants, virtual dynamic models of tomato development was presented using GREENLAB dual-scale automation (Dong *et al.*, 2010) and a parametric L-System (Li *et al.*, 2015; Xin *et al.*, April 16, 2014) at the organ level. To produce real models of tomato plants, a close range photogrammetric package, which is basically a correspondence based method, was used to generate 3D points of large tomato plants (Aguilar *et al.*, 2008). A correlation of R^2^ values of 0.75 between the measured volumes and manually-derived reference volumes was found, and the correlation of R^2^ values between the leaf area index (LAI) and manual volume was found to be 0.82. A combined SfM and MVS technique was carried out in 2015 (Rose *et al.*, 2015); the phenotypic traits, including leaf area, main stem height and convex hull, of the complete plant were estimated and compared to the reference data acquired by a laser scanner, and high R^2^ values were found, greater than 0.9. In addition, Nguyen built a system by integrating 5 stereo camera pairs, a structured light system, and software algorithms to model plants (Nguyen *et al.*, 2015), which was basically a correspondence-based approach. The results reached a recall of 0.97 and a precision of 0.89 for leaf detection and less than a 13-mm error for plant size, leaf size and internode distance. However, there are some shortcomings of the previous work, such as the lack of a real 3D model of tomato plants, the high cost of hardware systems, and the fact that it takes only certain growth stages into consideration. In other words, the potential of 3D phenotyping technologies has not been fully exploited on tomato plants over the full life cycle.

Thus, 4DPhenoMVS, a low-cost and open-access 3D tomato phenotyping and growth analysis pipeline using a 3D reconstruction point cloud based on multiview images, is proposed to address the abovementioned issues. In addition, with dynamic 3D traits, we investigate the environmental influence on tomato plant growth and yield in greenhouses.

## Materials and Methods

### Image acquisition and data analysis pipeline of 4DPhenoMVS

Considering the advantages of image-based reconstruction approaches, such as noninvasiveness, flexible data collection, and low cost, we present a spatial-temporal method called 4DPhenoMVS to reconstruct 3D point clouds of tomato plants; this method covers the full life cycle of tomato plants based on the SfM-MVS approach, uses multiple images captured by a consumer camera, and calculates 3D tomato phenotypic traits. The proposed 4DPhenoMVS consists of 8 modules, as shown in Fig. 1 and listed as follows: (1) Environmental monitoring. A distribution map of environmental differences is illustrated for tomato plant sample selection in Fig. 1A. (2) Multiple-image data collection covering the whole life cycle of tomato plants, as shown in Fig. 1B. (3) 3D point cloud generation through the combined SfM-MVS algorithms. (4) Alignment of point clouds regarding one tomato plant. For large tomato plants, image data are collected at two height levels, and the generated point clouds need to be aligned and registered together to produce a complete representation of the whole tomato plant. (5) Segmentation of stalk and leaf point clouds. (6) Skeletonization of stalk point clouds for structural phenotypic trait estimation. (7) Node detection on the skeletonized stem point clouds. Node detection is conducted on the skeletonized points; hence, the phenotypic traits, including node number, internode length, and stem length, are calculated automatically. (8) Structural phenotypic trait calculation and analysis. The phenotypic traits covering the whole life cycle of each tomato plant are calculated, as listed in Table 1.

**Figure 1.**
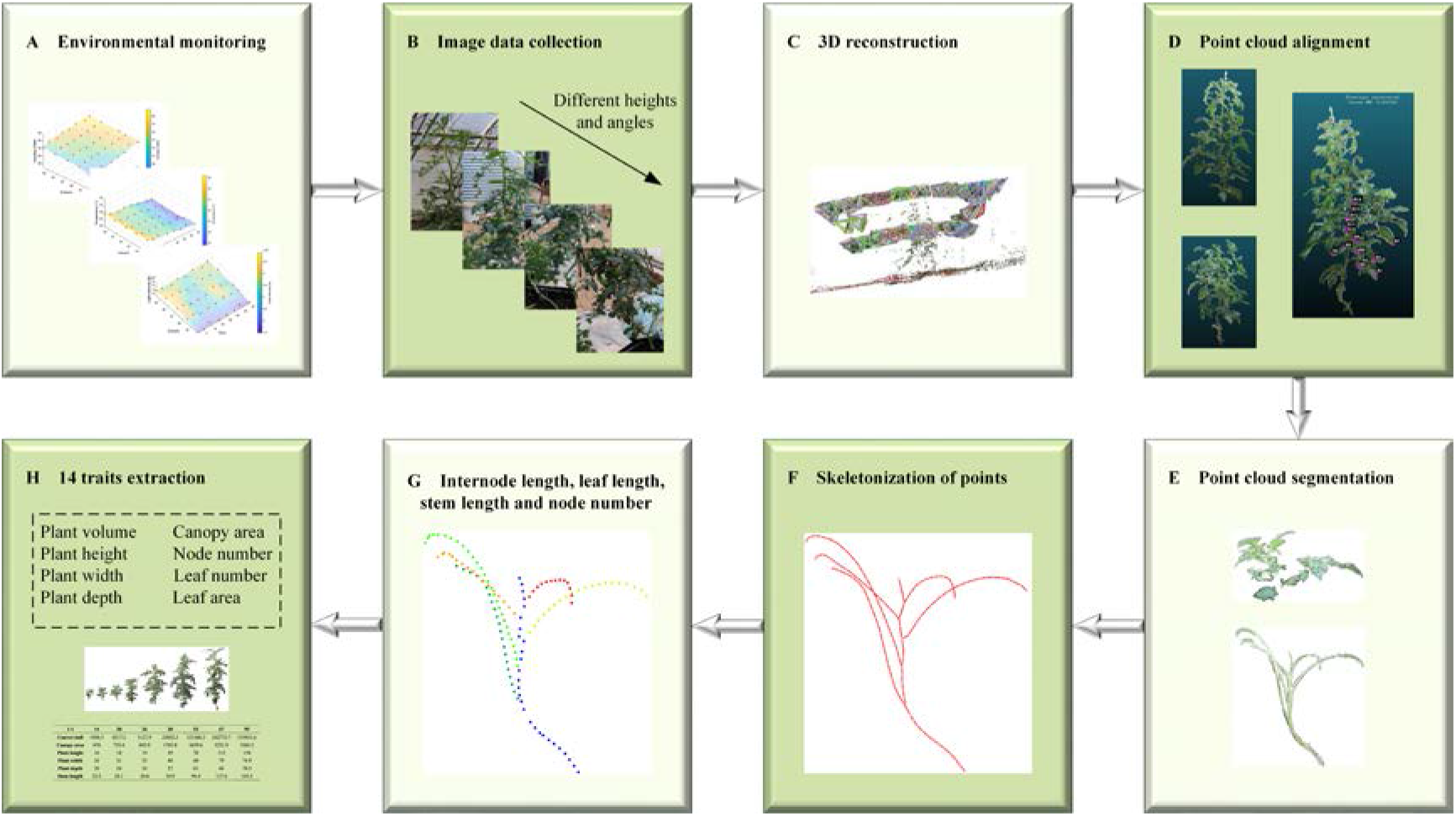
4DPhenoMVS: A structural phenotypic trait estimation and growth analysis pipeline for tomato plants, covering the whole life cycle. (A) Environmental monitoring to determine the target plants, (B) image data collection to obtain multiview image sequences, (C) 3D reconstruction from images to generate 3D point clouds, (D) point cloud alignment to output a complete tomato plant model, (E) stalk and leaf point cloud segmentation to separate stalk point clouds and leaf (except petiole) point clouds, (F) skeletonization of stalk point clouds, (G) phenotypic trait extraction and visualization based on the skeletonization results, and (H) extraction and analysis of 14 phenotypic traits.

**Table 1.** Phenotypic traits and their abbreviations.

| **Trait** | **Abbreviation** |
| --- | --- |
| Plant height: height of the minimum bounding box | PH |
| Plant width: width of the minimum bounding box | PW |
| Plant depth: depth of the minimum bounding box | PD |
| Plant volume: convex hull volume | PV |
| Stem length: sum of all internode lengths | SL |
| Canopy area: product of plant width and depth | CA |
| Primary ratio: ratio of plant height to plant width | RP |
| Secondary ratio: ratio of plant height to plant depth | RS |
| Internode length: Euclidean distance between two adjacent nodes | IL |
| Node number | NN |
| Leaf number | LN |
| Leaf length: sum of all the Euclidean distances between two ends | LL |
| Leaf width: width of the minimum bounding box of the leaf | LW |
| Total leaf area | TLA |

### Plant material and image data collection

Due to the superiorities of the image-based reconstruction method mentioned above, a consumer-cost digital camera (EOS 77D, Canon Corporation, Japan) was employed in our study. The tomato plants were planted in troughs in a form of soilless culture in a greenhouse that was approximately 400 m^2^. Tomato seedlings were transplanted into the greenhouse on 1 August 2020, which is located in Liaocheng City, Shandong Province, China (Fig. 2). The greenhouse was equipped with common management facilities, including reservoirs, drip irrigation and drainage components, draught fans, wet curtains, and vents. The reservoir and wet curtains were located at one end of the greenhouse, and the draught fan was located at the other end. There were two vents, upper and lower vents, allowing for air circulation. A total of 35 rows of plants were planted, with 55 plants in each row. The row spacing was 0.8 m, and the plant spacing was 0.16 m. The plant height was approximately 0.3~0.4 m, and the seedlings were hung 17 days after planting. The planting scenario is illustrated in Fig. 2.

**Figure 2.**
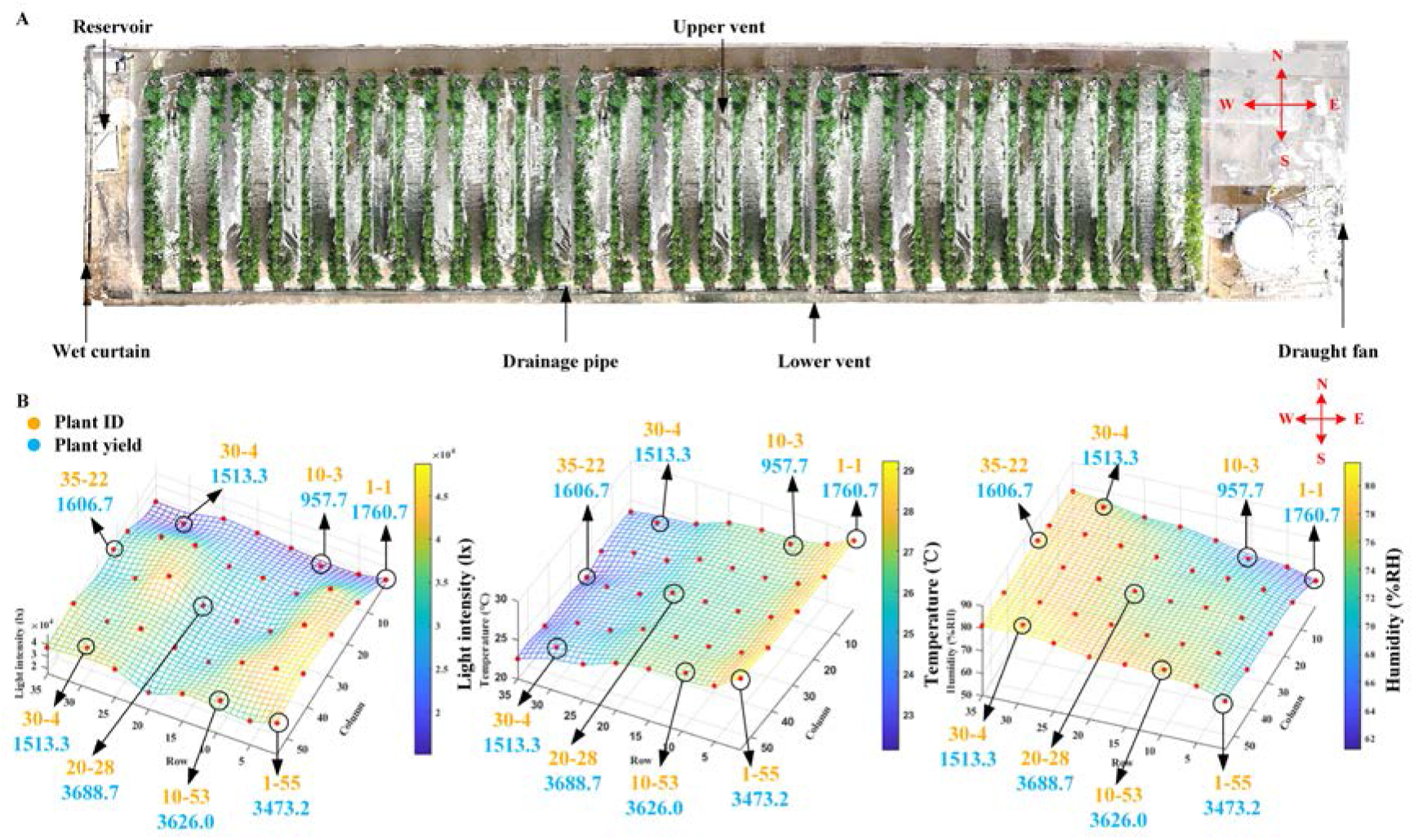
The plant scenario in the greenhouse. (A) Top view of the greenhouse and the locations of draught fans, reservoirs, wet curtains, drainage pipes, and vents. (B) The environmental parameters vary in different locations with light intensity changes (left), temperature changes (middle), and humidity changes (right). The yield for each individual tomato plant is also shown.

Ideally, the environmental conditions inside the greenhouse would be consistent; however, there were still some local microenvironment differences (microclimates) in the greenhouse. Hence, the local environmental parameters, including temperature, humidity, and light intensity, in the greenhouse were measured. The temperature and humidity were measured by a RENKE COS03 Temperature data logger (Shandong Renke Control Technology Co., Ltd, China), and the light intensity was measured by an EVERFINE PLA-20 plant lighting analyzer (EVERFINE Corporation, China). To avoid differences due to manual intervention, 40 sample locations of environmental factors were first monitored and recorded evenly in the greenhouse. The measurement work lasted for 8 consecutive days. The change in the environmental parameters for the 40 sample locations is illustrated in the 3D scatter plots shown in Fig. 2C.

As illustrated in Fig. 2A, draught fans were equipped at the east end of the greenhouse, and wet curtains were equipped at the west end. In addition, the upper vents were mounted in the northern area, and the lower vents and drainage pipes were set in the southern area. All the conditional parameters were measured under the same controlled conditions; i.e., the vents were closed, and the draught fans and wet curtains were open. Due to the effect of the wet curtain and the draught fan, the closer to the wet curtain, the higher the humidity was, and the closer to the draught fan, the higher the temperature was. Similarly, drainage troughs would lead to higher humidity in closer areas, and vents would lead to lower temperatures. In addition, occlusion of the shelter would cause lower light intensity. A closer look at Fig. 2C, 2D and 2E shows that gradual changes occurred for all three environmental parameters. Based on the discussion above, 8 tomato plants located in the 8 positions where environmental conditions obviously varied were selected as the target samples in this study, and they are illustrated by circles in Fig. 2C, 2D and 2E.

For convenience of description, the notation “plant i-j” is used to represent one plant, where i represents the number of the row in which the tomato plant is located, which ranges from 1 to 35, and j represents the number of the column in which the tomato plant is located, which ranges from 1 to 55. The selected target samples were as follows: plants 1-1, 1-55, 10-53, 30-4, 30-52, 35-22, 10-3, and 20-28.

For each plant, the user moved the camera around the tomato plant in a circular fashion, and approximately 100 images were taken for each tomato plant. Image collection started on the 14^th^ day after transplantation. When the tomato plants grew very quickly in the early stage, images were collected every 6 days. Since the growth rate of the tomato plants decreased in the later period, images were taken every 15 days until the tomato fruits were mature. As the plant grew, self-occlusion often occurred because the bottom parts could be occluded by the top parts. In this case, the digital camera needed to be set at different height levels, and two sets of images were collected in a circular fashion. The data collection scenario is illustrated in Fig. 3.

**Figure 3.**
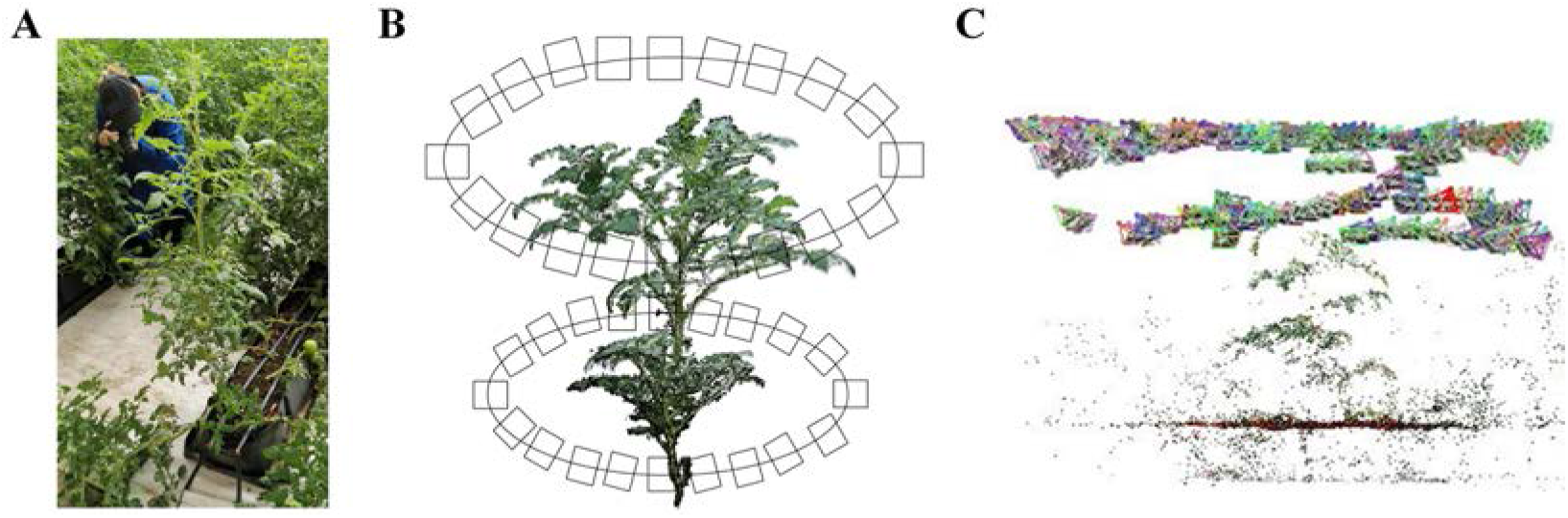
Data collection scenario and 3D reconstruction. (A) Photographing a tomato plant with the camera. (B) The path of the camera at two different height levels during image acquisition. (C) Reconstruction process of a tomato plant model.

### 3D Reconstruction of tomato plants by SfM

SfM is a photogrammetric range imaging technique for estimating 3D structures from image sequences. It takes multiple images as input and outputs the camera parameters for each image as well as the coarse 3D shape of the object; that is, it performs sparse reconstruction. Usually, key point feature detection algorithms, such as the scale-invariant feature transform (SIFT) image feature descriptor (Lowe, 2004)or variations of it, were implemented on the input images and later used for key point matching, and bundle block adjustment from the photogrammetry community was also introduced to estimate the accurate camera parameters for each image (Wu *et al.*, 2011). Through these procedures, the intrinsic and extrinsic parameters for each image were calculated properly, and the sparse points of tomato plants were also generated. The MVS algorithm then used the calibrated images to derive a very dense point cloud, which was color-coded using the original image data, by adopting epipolar geometry. The above procedures can be implemented with the free academic software VisualSFM (Wu, 2013; Wu *et al.*, 2011). In cases where two sets of point clouds for the upper and lower parts of one tomato plant were generated, the complete point cloud could be created by aligning the two sets of point clouds by locating 3 or more correspondences between them.

### *L*_*1*_-*medial* skeleton of the stalk point cloud

Minor manual work was involved to separate the leaf points and stalk points, as shown in Fig. 1E. Then, based on the stalk points, extracting a skeletal representation was an effective tool for geometric analysis and manipulation. In addition, a significant decrease in the time cost for the geometric analysis of the skeletal representation could be achieved compared with the analysis of the original point cloud. An *L*_1_-*medial* skeleton algorithm (Huang *et al.*, 2013) was demonstrated to be an effective method without prior assumptions and prior processing, which may include denoising, outlier removal, normal estimation, data completion, or mesh reconstruction. It adapted *L*_*1*_-*medians* locally to a point set representing a 3D shape, giving rise to a one-dimensional structure, which was seen as the localized center of the shape. Thus, this would be an appropriate skeleton algorithm for use on the reconstructed point cloud, which might be uneven and incomplete. Although competitive skeletal results can be derived, time efficiency is seriously affected by the data size of the input point cloud. During SfM-MVS reconstruction processing, many redundant points were produced, which would lead to low efficiency for phenotypic trait estimation. Therefore, the original point cloud was downsampled through an octree-based methodology to balance the trade-off between efficiency and performance. The downsampled point cloud was set as the input for the skeletonization process. Diagrams showing the original point cloud and the generated skeleton are provided in Fig. 1E and 1F.

### Structural phenotypic trait estimation

Structural phenotypic traits represent the morphological characteristics of tomato plants, such as plant volume, plant height, canopy height, canopy area, node number and internode length, and the definitions and abbreviations of the phenotypic traits are illustrated in Table 1.

#### Plant height

Plant height is defined as the vertical distance from the ground to the highest point of the plant in its natural state, which can be represented by the height of the minimum bounding box, *H*_B_.

#### Plant volume

Plant volume can be represented by the volume of the convex hull and the minimum bounding box (MMB). The convex hull is the smallest convex set that contains all the points, while the MMB is defined as the minimum box that contains all the points of the tomato plant. Fig. 4 shows the 3D tomato plant model enclosed by the convex hull and MMB.

**Figure 4.**
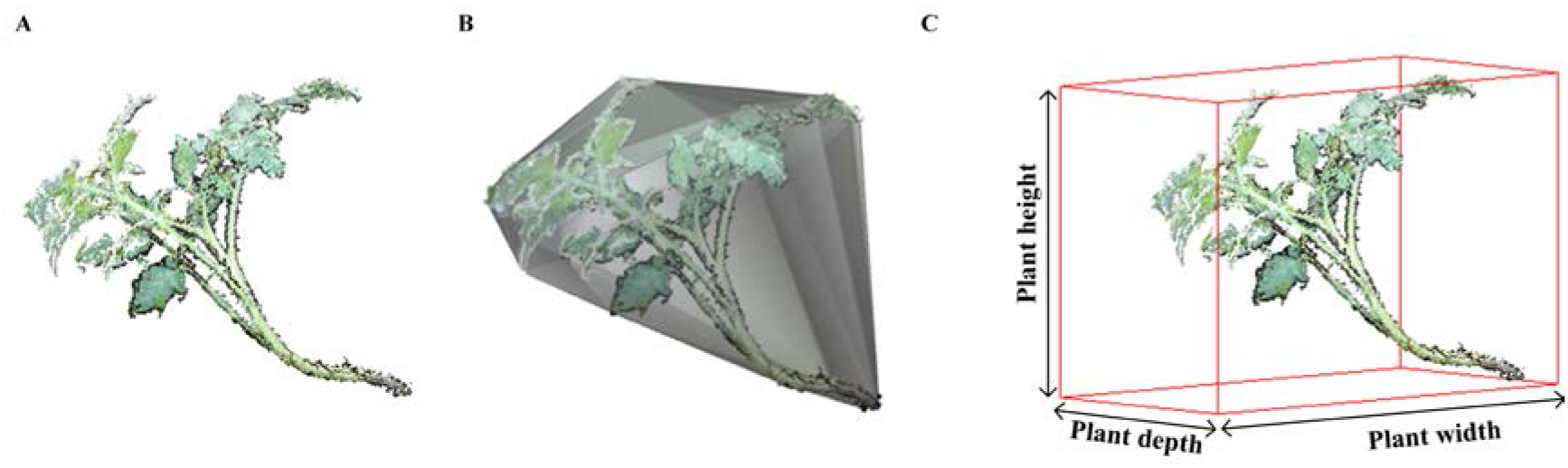
Tomato plant volume estimation. (A) Original point cloud for one tomato plant. (B) Convex hull of the point cloud used for volume calculation. (C) MMB of the point cloud for one tomato plant.

#### Plant volume

Plant volume can be represented by the volume of convex-hull, and the MMB (minimum bounding box), respectively. The convex-hull is the smallest convex set that contains all the points, while the MMB (minimum bounding box) was defined as the minimum box that contains all the points of tomato plants. Fig. 4 shows the 3D tomato plant model enclosed by the convex hull and MMB.

#### Stem length

Stem length is defined as the sum of all internode lengths.

#### Canopy area

The canopy area is defined as the multiplication of the plant width and plant depth.

#### Node number and internode length

The main stem of the tomato plant is not typically straight, which will make the measurement of plant height difficult. A series of nodes are located around the main stem. The internode length is defined as the Euclidean distance between two adjacent nodes. The length together with the number of internodes determines the length of the main stem. Hence, an alternative strategy for stem length calculation is adopted in the study. The skeleton of the stalk is already generated on the point cloud, and it depicts the morphological structure. The nodes located in the main stem can be found by using a suitable search strategy. In this way, the internode length, representing the distance between two nodes along the main stem, can be calculated, and the stem length can be determined by summing all the internode lengths.

#### Number of leaves, leaf length and leaf area

The number of leaves denotes the total number of leaves in the plant, and it indicates the tomato plant’s physiological age; thus, it is also involved in phenotypic trait estimation. In the early growth period of the tomato plant, it is identical to the number of nodes; however, it is different in the late growth period due to the pruning of old leaves. Leaf length measures the length of the leaf in 3D space, and it is estimated by summing all the Euclidean distances between two leaf ends. One end is the starting point of the petiole located at the main stem, and the other end is located at the end of the leaf. Leaf area is the most closely related and variable factor of yield. It measures the surface area of leaves in 3D space. To estimate one leaf area accurately, a mesh model for the 3D points representing this leaf is generated, and the sum of the area of all the meshes is the leaf area.

## Results

### Dynamic 3D reconstructed point cloud of tomato plants

For each individual tomato plant, one group of images was collected during the earlier growth stage, while two sets of images, for the upper part and lower part, were acquired once occlusion occurred during the later growth stage. Alignment was implemented to produce a complete point cloud with respect to the whole plant, as shown in Fig. 5A–5C. Time series 3D models of tomato plants covering the full growth period were created with high-quality and rich color information, and the structural changes of tomato plants during the different growth and development stages were also visualized, as shown in Fig. 5D. After removing the leaf points manually, only the stalk points were retained, and the skeletonization algorithm was implemented on the downsampled stalk points to output skeletonized stalks with nodes. Fig. 5E shows a side view of the skeletonized results. Node detection was conducted in an automatic way for node number and internode length calculation. (The source code and detailed implementation are described in Supplementary Note. S1)

**Figure 5.**
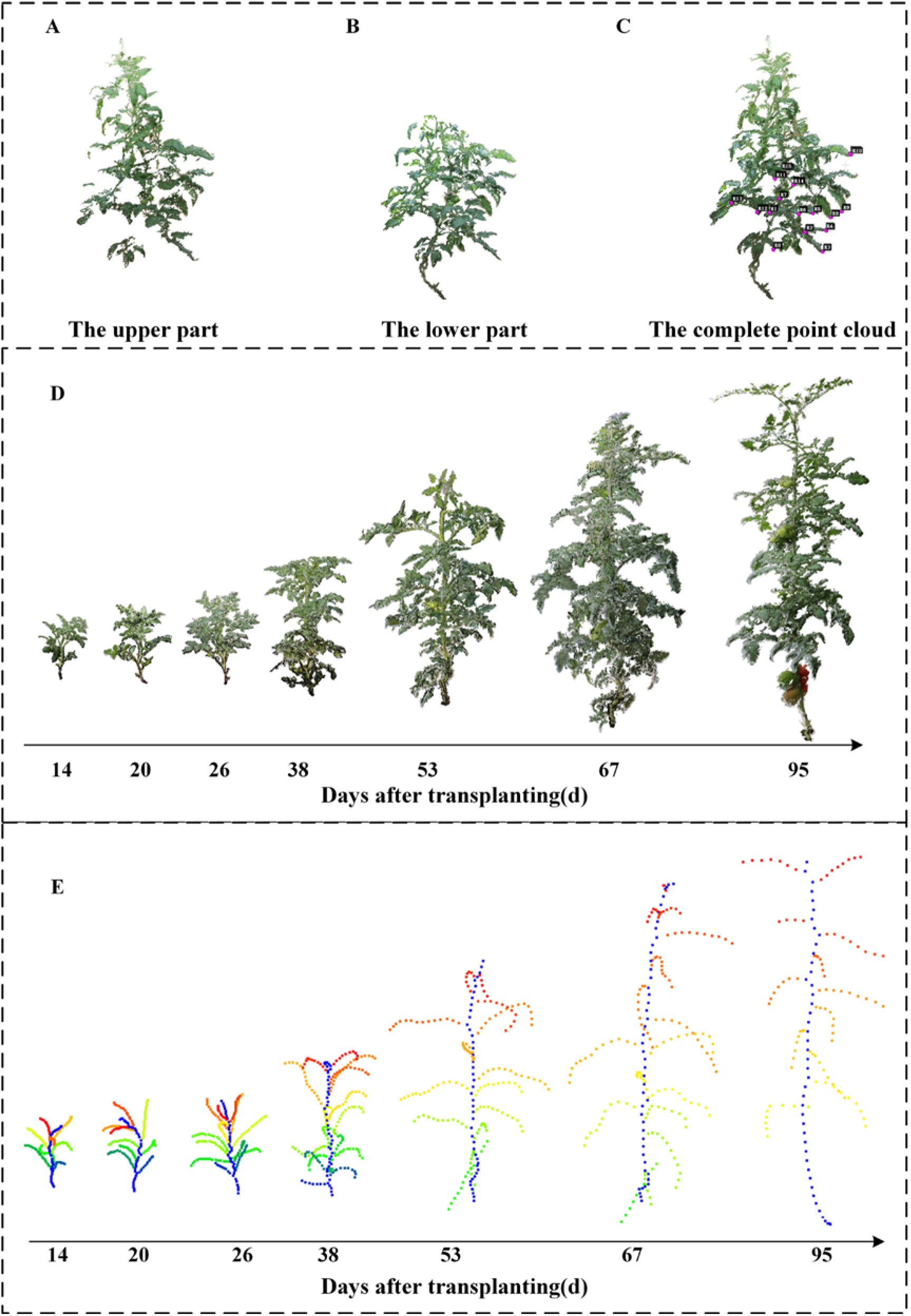
Visualization of reconstruction and stalk skeletonization. Point clouds for the upper part (A) and lower part (B) of one tomato plant and (C) a complete tomato plant model after alignment. (D) All growth stages of one tomato plant at 7 time points. (E) The stalk skeletonization results of one tomato plant at 7 time points.

### Performance evaluation on structural phenotypic traits

Manual measurements of phenotypic traits were also conducted on tomato plants to demonstrate the accuracy of the algorithm proposed in this research. The comparative results are shown in Fig. 6. Plant height and internode length were recorded by using a measuring tape, and the node number was also counted and recorded. The results showed that the R^2^ values of plant height, stem length, internode length, and leaf number were 0.96, 0.96, 0.83 and 0.76, respectively. The mean absolute percentage error (MAPE, defined in Equation (1)) of plant height, stem length, internode length, and leaf number were 17.23%, 13.89%, 20.62% and 8.20%, respectively

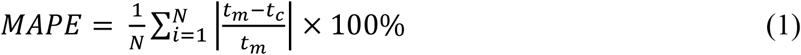

where *N* denotes the number of times the summation iteration occurs, *t*_*m*_ denotes the measurement value, and *t*_*c*_ denotes the value calculated from the 3D models.

**Figure 6.**
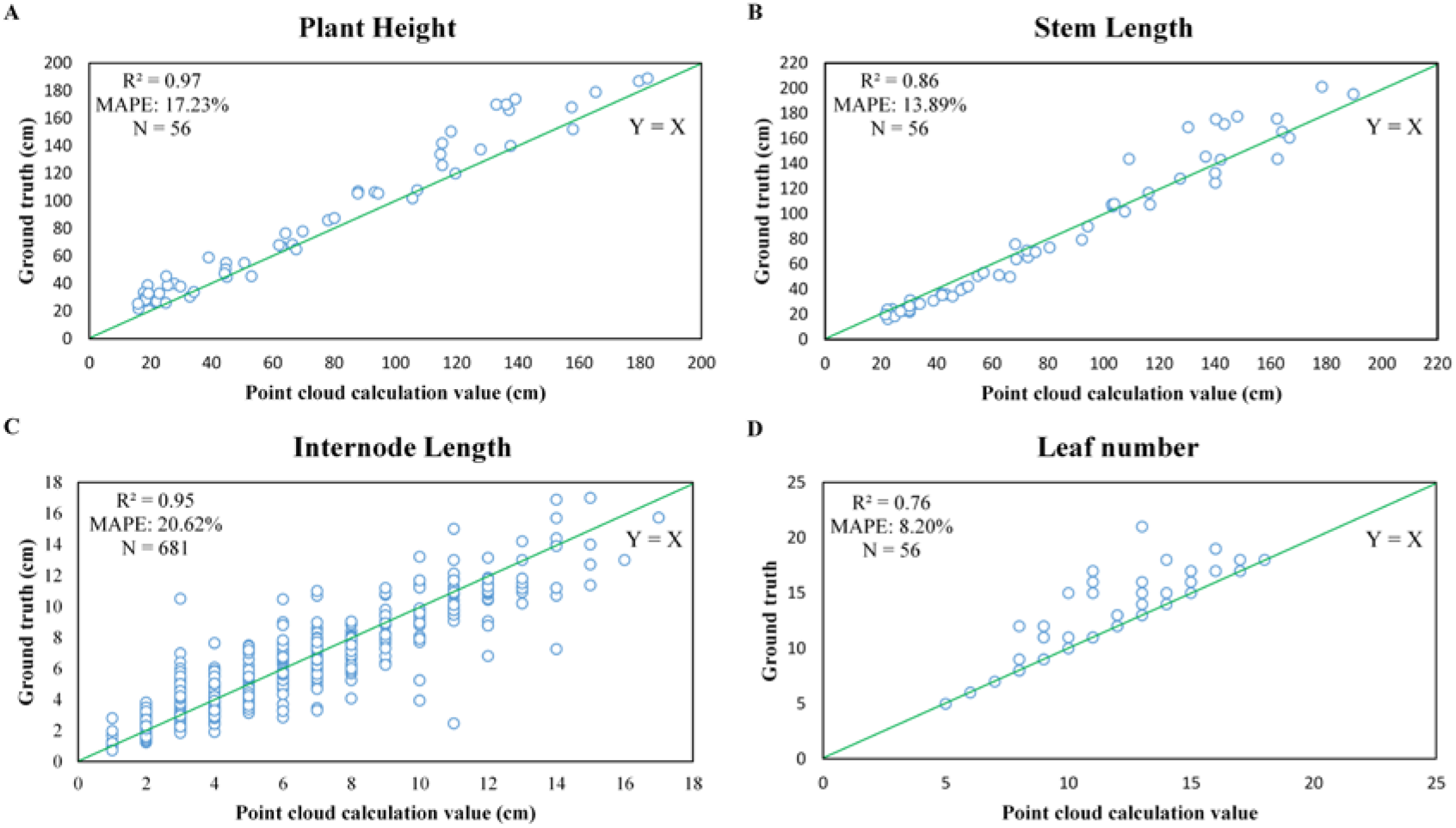
The comparison of phenotypic trait parameters between the manual measurements and estimation based on the generated 3D models. (A) Plant height, (B) stem length, (C) internode length, and (D) leaf number.

### Growth variation with different environments

To investigate the environmental influence on the growth of tomato plants, the dynamic growth variation in different environments was measured and validated. As shown in Fig. 7, the three environmental factors showed various influences on the growth of the tomato plants planted in the greenhouse. Tomato plants located in areas with higher intensity and moderate temperature and humidity show longer stem lengths than plants with lower light intensity, such as plants 1-55, 10-53, and 20-28. Plants 10-3 and 30-4, located at the other end of the greenhouse, show shorter stem lengths and lower growth rates of stem length. Considering the influence of temperature, the tomato plants (plants 1-55, 10-53, and 20-28) in the higher-temperature area show relatively large plant height, while plants 30-4, 30-52, and 35-22 show lower plant height. Two rows of plants (rows 30 and 35) were located at the west end of the greenhouse, where a reservoir and wet curtains were equipped; hence, the temperature was lower than that on the other side. Regarding humidity, the two plants located in the 30^th^ row with high humidity, plants 30-4 and 30-52, showed lower plant volumes, while the plants located in the area with lower humidity showed higher volumes, as shown in Fig. 7E and 7F.

**Figure 7.**
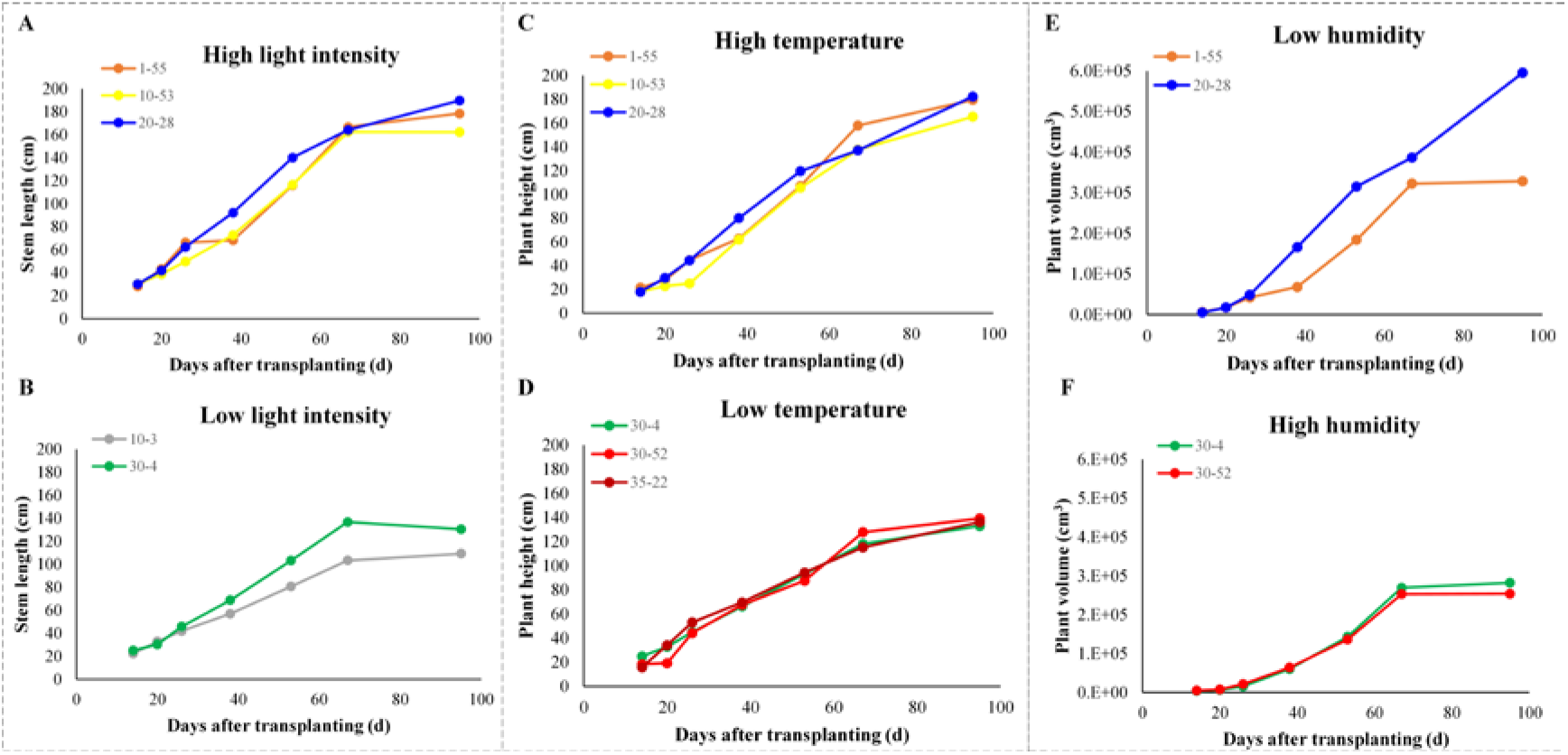
Variation in phenotypic traits influenced by different light intensities, temperatures, and humidities. (A-B) Influence of light intensity variation on stem length, (C-D) influence of temperature variation on plant height, (E-F) influence of humidity variation on plant volume.

In addition, tomato fruit weight for each individual plant was measured to further investigate the environmental influence. As shown in Fig 2C–2E, higher yields occurred in plants 1-55, 10-53, and 20-28, while plants 10-3 and 30-4 showed lower yields. Thus, by testing the eight samples in the greenhouse, it was determined that higher yield can be expected for plants with higher light intensity and moderate temperature and humidity.

### Measuring efficiency evaluation

In this study, a consumer digital camera was used to collect image data for each individual tomato plant. While the plant was in the seedling stage, approximately 100 images were taken, and this took approximately 15 minutes; in the later stage, two sets of images were acquired, and approximately half an hour was needed. These procedures were all implemented on a Windows 10 operating system, and the central processing unit (CPU) was an Intel Core i7-8700 (16 GB of random-access memory). SfM was implemented by VisualSFM, and approximately 60 minutes were needed to generate one point cloud for one tomato plant. Plant volume was represented by the convex hull of the generated 3D points, and approximately 1 minute was needed to calculate the volume of a single plant at 7 time points. Plant height, plant width and plant depth were represented by the minimum bounding box of the generated 3D points, and determining these values took approximately 20 seconds. Leaf points were separated from stalk points by minor manual work, which took approximately 2 minutes. After that, the *L*_1_ *median* skeletonization algorithm was carried out on the downsampled stalk points, and it took approximately 1-2 minutes to process 10k points. Node detection was conducted on the skeletonized stalk points to calculate internode length automatically (the source code for node detection can be downloaded as explained in Supplementary Note. S1). Thus, the total time consumption for extracting all the morphological traits for one tomato plant was 80 minutes.

Among all the procedures, the 3D reconstruction process took most of the time, which is mainly due to the structural complexity of tomato plants, as can be seen from the visualization results. Higher reconstruction efficiency would be achieved with a more advanced computer configuration. In addition, the proposed pipeline has another significant advantage: all the node detection and internode length calculation procedures regarding the stalk of one individual tomato plant at different time points are implemented in a fully automated way. The phenotypic trait calculation regarding leaves follows the same pipeline. In other words, it is not necessary to build the correspondences for certain organs, such as nodes and leaves, at different growth stages in order to investigate the temporal trait changes regarding one plant organ.

## Discussion

### Reconstruction strategy

To date, the development of 3D reconstruction technology based on multiple images has promoted a wide range of applications for this technology in 3D plant phenotyping because of its advantages as a non-contact, low-cost and high-precision method. Fixed cameras and moving cameras are the two main scenarios used for plant reconstruction. In the fixed-camera scenario, the advantage of this approach is its high time efficiency for 3D reconstruction, and fewer images are required as input data in this case than in the moving-camera scenario. However, some notable types of defects cannot be prevented by this solution. The prerequisite for fixed-camera reconstruction is to calculate the accurate position and orientation parameters of all cameras, i.e., the intrinsic and extrinsic parameters of the cameras, which are determined through a calibration algorithm. This increases the complexity of the approach, especially for researchers who do not have computer vision experience and backgrounds. Both the carving-based and correspondence-based reconstruction methods are involved in the calibration procedures; thus, other accessibility settings, such as a checkerboard (Zhu *et al.*, 2020) or a cube with checkerboard patterns (Das Choudhury *et al.*, 2020), were used for the calibration procedure. In addition, multiple image data collection was completed with the assistance of a turntable or stand (Liu *et al.*, 2017), an electronic rotary (Li, 2019) and other auxiliary facilities. Second, the experiment was limited to specific indoor scenes because of the utilization of fixed cameras and other accessibility settings.

Regarding the investigation of the environmental influence on tomato plants, all the environmental conditions should remain the same, and the tomato plants were fixed at the same locations during all growth stages. Hence, moving-camera solutions were more applicable in this study. In contrast to the fixed-camera scenario, complicated calibration procedures were avoided, and the tomato plants could stay in their original environmental conditions without human intervention.

### Effect of environmental variations

It has been demonstrated that environmental differences influence the growth of plants. No significant growth difference regarding the total plant area, plant volume, stem length or canopy area was shown in the early periods, while a great difference was shown in the late growth stages, especially during the fruit growth periods.

Differences in light intensity, temperature, and humidity can lead to differences in plant growth. In the northern part of the greenhouse, the light intensity was weak due to the shelter. Wet curtains led to higher humidity, and draught fans and vents led to lower temperatures. In the same row, the difference in temperature and humidity was very small. Interestingly, the plant growth on the south side was better than the plant growth on the north side. For example, the overall growth of plants 1-55, 10-53 and 30-52 was better than that of plants 1-1, 10-3 and 30-4, respectively. This may indicate that light intensity has a greater impact on plant growth. It should be noted that the three environmental parameters in the central area were not the highest or lowest, but the yield of plant 20-28 was the highest. The reason was mainly because the environmental changes in the central area were not as drastic as those on the east or west sides when performing normal switching operations.

### The environmental influence on the yield

To investigate the environmental influence on the yield, the fruit number and fruit weight were also measured manually at the mature stage. As shown in Fig. 2C–2E, the yields of plants 1-55, 10-53, and 20-28 were higher than those of the other 5 plants, indicating that higher light intensity with moderate temperature and humidity will contribute to better yield. In other words, slight differences in environmental changes, especially in light intensity, caused significant differences in individual plant growth and ultimately affected the yields of tomato plants.

### Outlook and perspectives

The application of the proposed low-cost and open-access pipeline has been demonstrated on tomato plants covering all growth stages. To further exploit the application of the proposed 4DPhenoMVS, two aspects need to be considered in future research: (1) The pipelines could be applied to other horticultural crops, such as potato and lettuce plants. (2) To exploit the relations between plant growth and environmental conditions, further experiments could be designed, for example, using a controlled light source to monitor the plant growth, such as leaf color or the rate of photosynthesis.

## Conclusion

In this work, a low-cost and open-access temporal 3D phenotyping pipeline, 4DPhenoMVS, was proposed. Based on the analysis of environmental differences in a greenhouse, eight tomato plants were chosen to generate 3D point clouds. The 3D reconstructed models could show the morphological structure covering the whole life cycle. In addition, simple skeletons of stalks were produced on the point clouds to perform accurate phenotypic trait calculations. Fourteen phenotypic traits were calculated, and the R^2^ values regarding stem length, plant height, and internode length were more than 0.8. In addition, slight environmental differences in the greenhouse had different influences on tomato plant growth, and the yield difference also demonstrated this. Specifically, stronger light intensity with moderate humidity and temperature contributed to higher yield. Overall, the proposed low-cost and open-access phenotyping pipeline could benefit tomato breeding, cultivation research, and functional genomics in the future.

## Supplementary Data

Supplementary data are available at *JXB* online.

**Note. S1.** The instruction of 4DPhenoMVS.

**Video.S1.** Operating procedure for 4DPhenoMVS.

**Video.S2.** The whole grow stages of plant 1-1 at 7 time points.

**Video.S3.** The whole grow stages of plant 1-55 at 7 time points.

**Video.S4.** The whole grow stages of plant 10-3 at 7 time points.

**Video.S5.** The whole grow stages of plant 10-53 at 7 time points.

**Video.S6.** The whole grow stages of plant 20-28 at 7 time points.

**Video.S7.** The whole grow stages of plant 30-4 at 7 time points.

**Video.S8.** The whole grow stages of plant 30-52 at 7 time points.

**Video.S9.** The whole grow stages of plant 35-22 at 7 time points.

## Acknowledgements

This work was supported by grants from the National Natural Science Foundation of China (31770397), Major science and technology projects in Hubei Province, Fundamental Research Funds for the Central Universities (2662020ZKPY017, 2021ZKPY006), Cooperative funding between Huazhong Agricultural University and Shenzhen Institute of agricultural genomics (SZYJY2021005, SZYJY2021007). We thanked Harvest-Code Technology (Nanjing) Ltd. provided the materials and experimental resources.

## Data Availability

All the phenotypic data and images can be viewed and downloaded via the link (http://plantphenomics.hzau.edu.cn/download_checkiflogin_en.action).

